# The Effects of Normalization, Transformation, and Rarefaction on Clustering of OTU Abundance

**DOI:** 10.1101/259325

**Authors:** David Molik, Michael Pfrender, Scott Emrich

## Introduction

Metagenomic clustering presents a unique opportunity to associate and understand communities. Working with Operational Taxonomic Units (OTUs), however, often requires a strategy for handling OTUs that may be over or under represented in a given sample, which is thought of as “erroneous”. PCR amplification, for example, is known to sometimes non-linearly over-represent more common species *Gonzalez et al. 2012*. Strategies dealing with this bias include Normalization, Rarefaction, and Log Transformation. Here, we examine how methods to handle potential outlier observations affect *de novo* estimation of groups using both clustering and matrix factorization methods.

While log transformations may affect parametric tests *O’Hara and Kotze 2010* our primary interest is in clustering, where relative abundances among samples could affect the derived clusters *Weiss et al. 2017*. We examine similarity between the adjusted OTU tables and the original ones to determine if relationships are preserved after post-processing.

We consider four methods: Bray-Curtis Similarity, *de novo* determined clusters by K-means and PCA, determining groups using Non-Negative Matrix Factorization (NMF). Our goal is to adjust for potential over or under representation while keeping realtive sample abudnaces as similar as possible, a problem that finds analogy in Single Cell sequencing *Vallejos et al. 2017*. We find that Rarefaction and then Transformation have higher mantel statistics, as a matrix correlation *Legendre and Legendre 1998*, in relation to an unmodified OTU table than a Normalized OTU table, and this same observation applies also to K-means. Only for NMF does Log Transformation results appear similar to an unedited table.

## Methods

We estimate a Bray-Curtis distance matrix from each of four OTU tables *Fig. 1* and compare these matrices with a mantel statistic *Fig. 2* to see if Normalization, Rarefaction, or Log Transformation disrupts groups. Data was chosen that had three distinct sampling locations and three distinct OTU profiles *Crits-Christoph et al. 2013*. We used k=3 for K-means and NMF, which is described below.

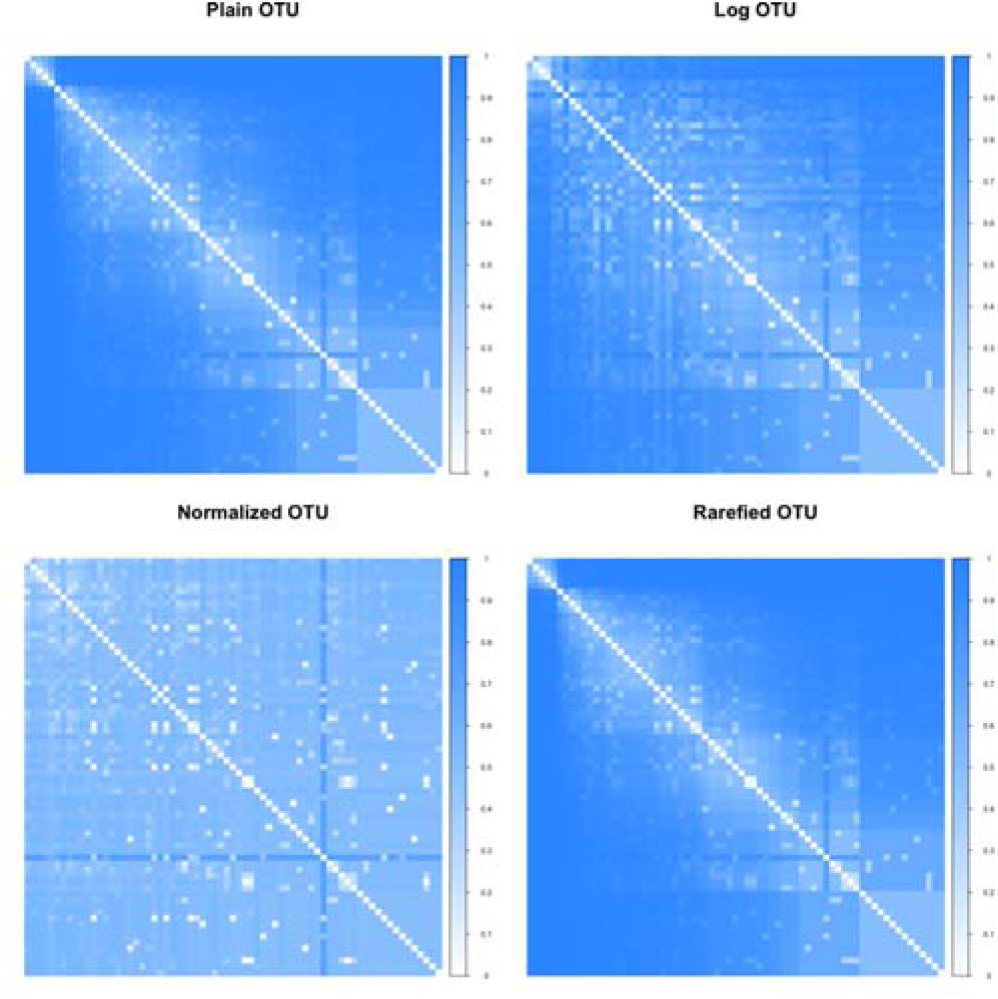

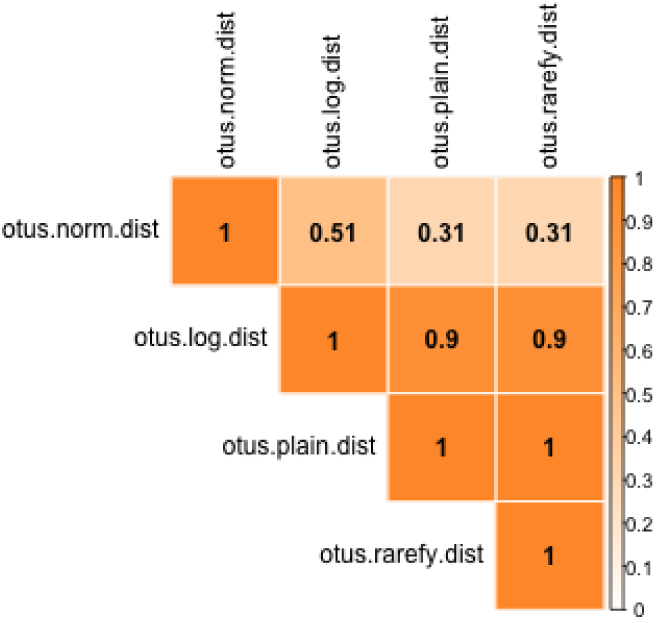

### K-means, PCA of Abundance

Using OTU tables we generate *de novo* clusters with K-means and then compare these clusters using a Jaccard-driven similarity *Fig. 3*. We test these clusters using a silhouette plot, which shows the distance of each sample to the closest other cluster. The clusters are plotted with a PCA on abundance, using the DC algorithm *Fig. 4*. Rand Similarities show the same relationships between OTU tables, and when a Wilcoxon signed-rank test is conducted between all tables, log transform is seen to have the highest statistic to the un-edited OTU table *Molik 2017*.

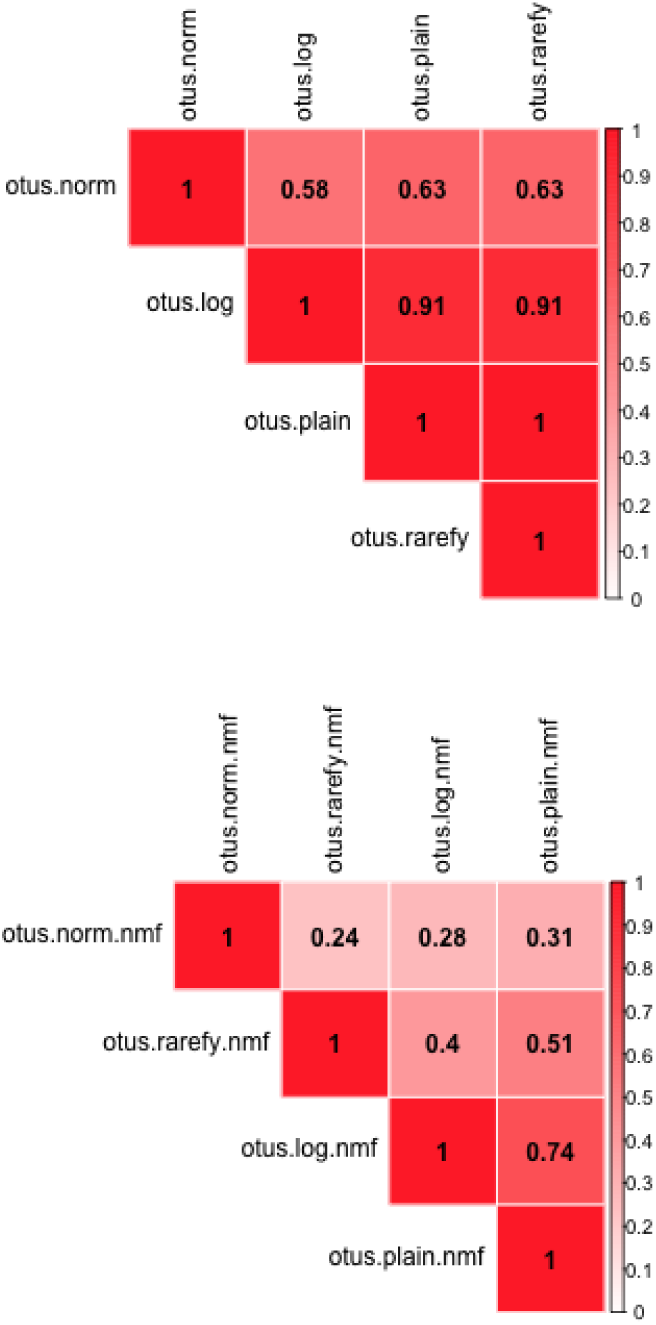

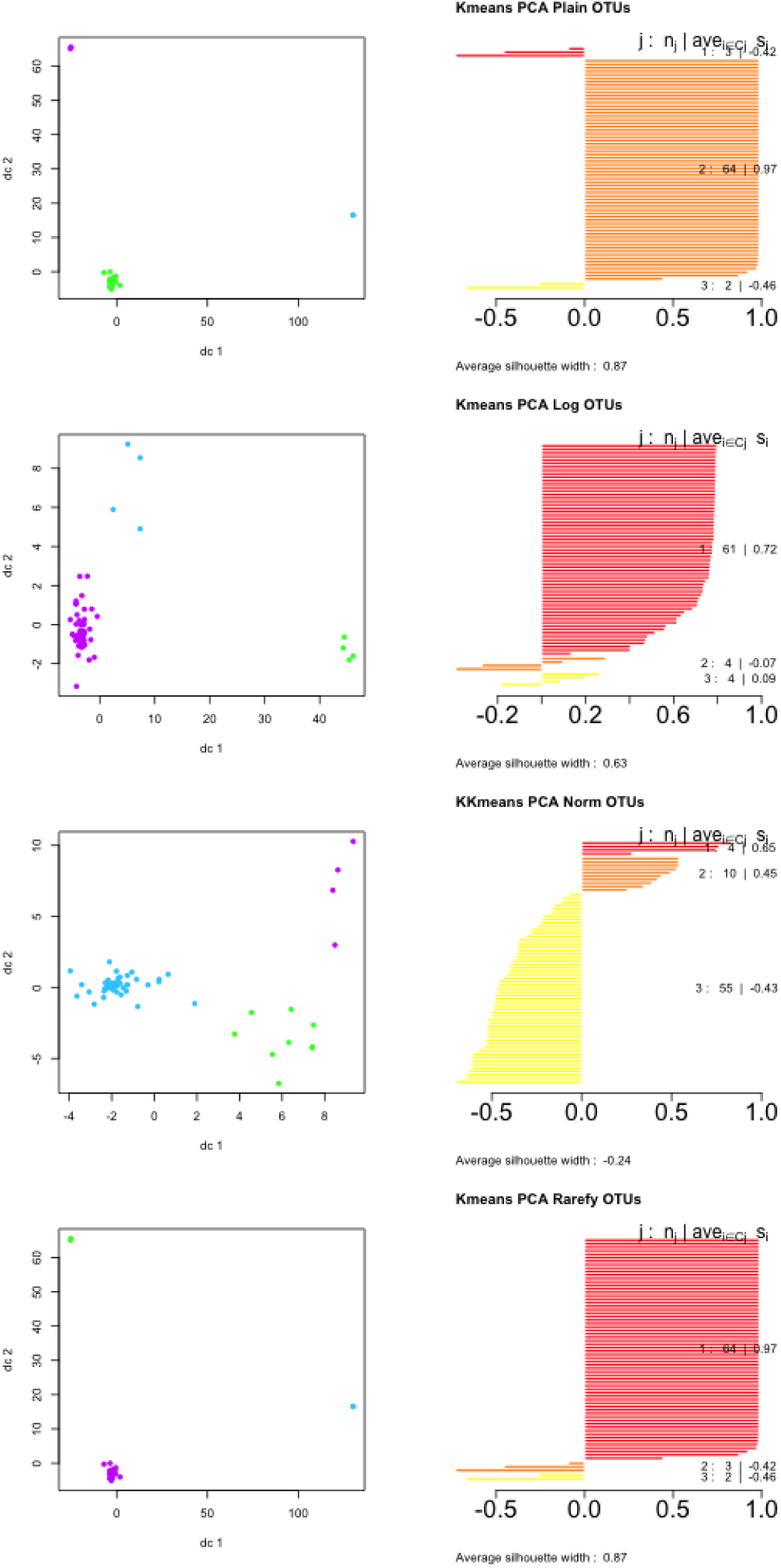

### Non-negative Matrix Factorization

Using the OTU tables we generate *de novo* clusters in NMF and compare these clusters with Jaccard Similarity *Fig. 3*. NMF groups had higher silhouette coefficients than their K-means counterparts *Fig. 5*. All groups had an above.8 silhouette coefficient.

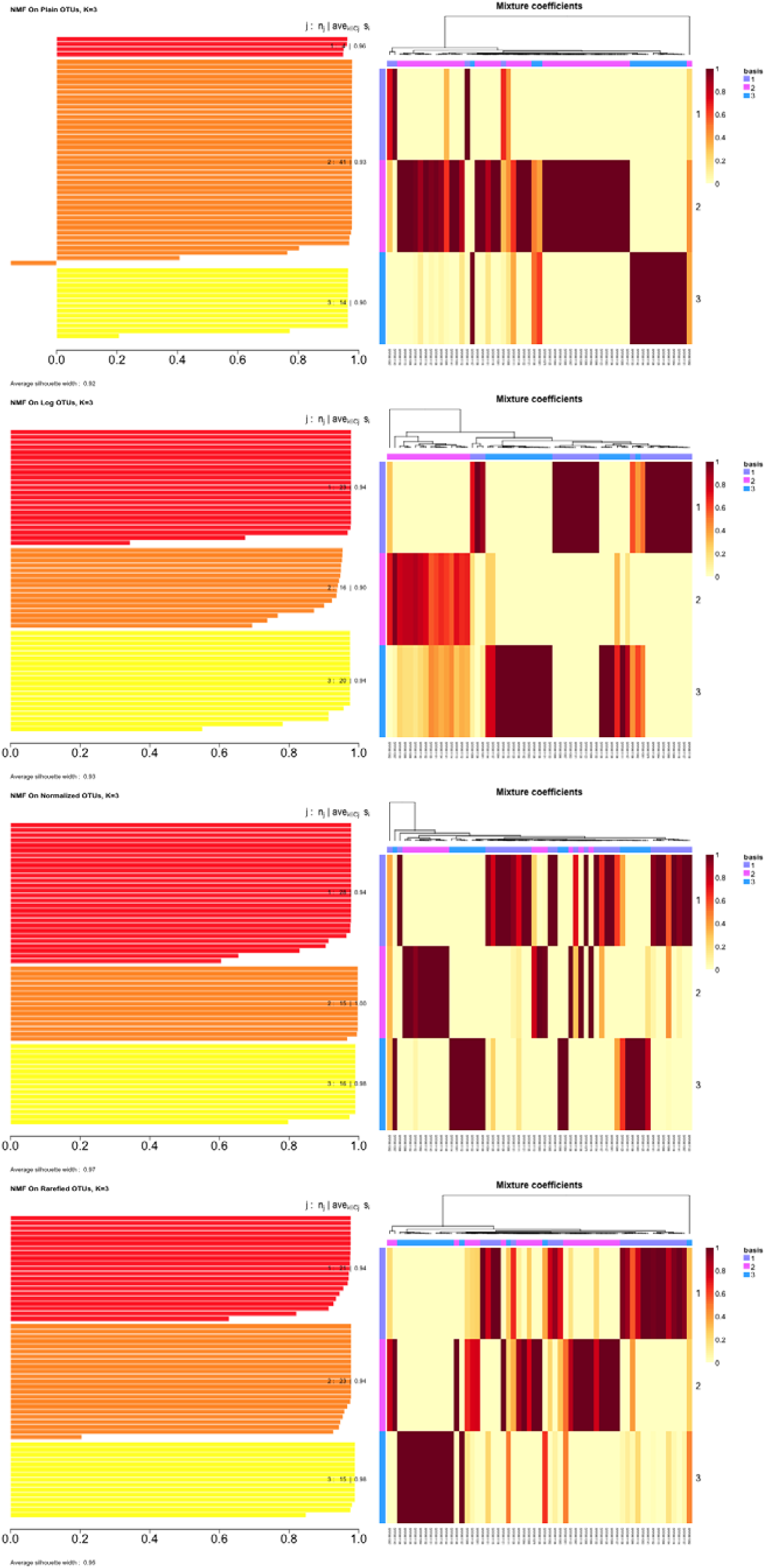

## Discussion

Distance matrices are used in hierarchical clustering. In Figure 2 we can observe that “normalized” is a fairly distinct distance matrix, and has been shown to cause a substantial difference in OTU composition *12*. These differences appear using mantel statistic assessment, as well as in K-means analysis and in NMF clustering. Similarly, the expectation in Rarefaction is that relative abundances will change according to the amount of rarefaction; a presence absence OTU table will have different sample to sample relationships than a slightly scaled down OTU table. We look at the later and, as expected, our rarefied table is shown to be more similar based on a mantel test on distance matrices, as well as in PCA/K-means assessments.

The optimal method for handling outliers and erroneous abundances depends on data and application *12*. In clustering and dissimilarity both rarification and log transformation seems to produce results that approximate the original OTU table; however, formatrix factorization only log transformation maintains original OTU table structure.

## Data and Code Availability

Source sequence data came from National Center for Biotechnology Information Sequence Read Archive SRA accession number SRA091062 *Crits-Christoph et al. 2013*. Analysis was conducted in R and is available at https://github.com/status-five/The-Effects-of-Normalization-Transformation-and-Rarefication-on-Clustering-on-OTU-Abundance, the DOI:10.5281/zenodo.896927 version was used *Molik 2017*. We used the R vegan package *Dixon 2003* for log transformation, normalization, and rarefaction of the OTU table. The NMF pacage was used for applying NMF *Gaujoux and Seoighe 2010*. The Corrplot R pacakge was used for matrices visualization *V. Simko 2016*. The fpc R package was used for PCA analysis and visualization *Hennig 2015*. The clusteval package was used for determining Jaccard similarities between clusters *Ramey 2012*.

